# Chirality-dependent anti-inflammatory effect of glutathione after spinal cord injury

**DOI:** 10.1101/2020.11.11.377721

**Authors:** Seong Jun Kim, Wan-Kyu Ko, Gong Ho Han, Daye Lee, Yuhan Lee, Seung Hun Sheen, Je Beom Hong, Seil Sohn

## Abstract

Neuroinflammation forms a glial scar following a spinal cord injury (SCI). The injured axon cannot regenerate across the scar, suggesting permanent paraplegia. In this study, we report that d-chiral glutathione (D-GSH) suppresses the inflammatory response after SCI and leads to axon regeneration of the injured spinal cord to a greater extent than l-chiral glutathione (L-GSH). After SCI, axon regrowth in D-GSH-treated rats was significantly increased compared to that in L-GSH-treated rats (^***^*p* < 0.001). Secondary damage and motor function were significantly improved in D-GSH-treated rats compared to those outcomes in L-GSH-treated rats (^**^*p* < 0.01). Moreover, D-GSH significantly decreased pro-inflammatory cytokines and glial scar via inhibition of the mitogen-activated protein kinase (MAPK) signaling pathway compared to L-GSH (^***^*p* < 0.001). In primary cultured macrophages, we found that D-GSH undergoes more intracellular interaction with activated macrophages than L-GSH (^***^*p* < 0.001). These findings reveal a potential new regenerative function of chiral GSH in SCI and suggest that chiral GSH has therapeutic potential as a treatment of other diseases.

## Introduction

Approximately 17,730 new cases of spinal cord injuries (SCI) occur for every year in the United States alone (Jain *et al*, 2015). SCI triggers a complex local inflammatory response, which is a critical pathophysiological process related to secondary damage after the initial SCI. Inflammation is a major feature of secondary damage and is excessively induced at a very early stage after an acute injury (Ko *et al*, 2019; Rust & Kaiser, 2017). High doses of methylprednisolone, a corticosteroid anti-inflammatory drug, have been used worldwide to treat acute SCI patients (Kim *et al*, 2017). However, there has been considerable debate over its use due to its adverse side effects (Qian *et al*, 2005; Sambrook *et al*, 1984; Topal *et al*, 2006).

Chirality is a unique property of molecules with identical elemental compositions but that are non-superimposable (Hu *et al*, 2019). Materials in the environment have one of two naturally selected enantiomeric forms. For example, amino acids in proteins show mostly left-handedness (L-form), whereas glucose in DNA and RNA show right-handedness (D-form). In biological systems, the chiral property can show entirely different functions due to chiral-specific interactions (Green *et al*, 2016; Yeom *et al*, 2020).

Glutathione (GSH) is a thiol-containing tripeptide that consists of cysteine, glutamic acid, and glycine. GSH is the most abundant antioxidant in living organisms (Bansal & Simon, 2018). Previous studies have reported that GSH induces the neuronal differentiation of rat stromal cells and improves functional recovery after SCI (Guizar-Sahagun *et al*, 2005; Sagara & Makino, 2008). However, these studies were only based on the L-chirality of GSH (L-GSH). To the best of our knowledge, the D-chirality of GSH (D-GSH) has not been investigated as to whether it affects SCI outcomes. Therefore, in this study, we investigated the different chiral effects of GSHs after SCI.

## Results and discussion

### Characterization of chiral GSHs

As shown in Figs 1a and b, chiral GSHs have completely identical compositions and are non-superimposable structures. To investigate the optical activity of chiral GSHs, we dissolved L- or D-GSH in distilled water and measured the optical activity using circular dichroism (CD) spectroscopy. As a result, L-GSH showed a strong negative peak, whereas D-GSH showed a strong positive signal at around 190 nm (Fig 1c). This indicates symmetrical CD signals and a distinct a Cotton effect at around 190 nm (Lowry, 1933). Litman and Schellman reported that the optical rotatory dispersion of peptides was correlated with the conformational and structural features of a molecule (Litman & Schellman, 1965). Taken together, we confirmed that L- and D-GSH had opposite chirality.

**Figure 1.**
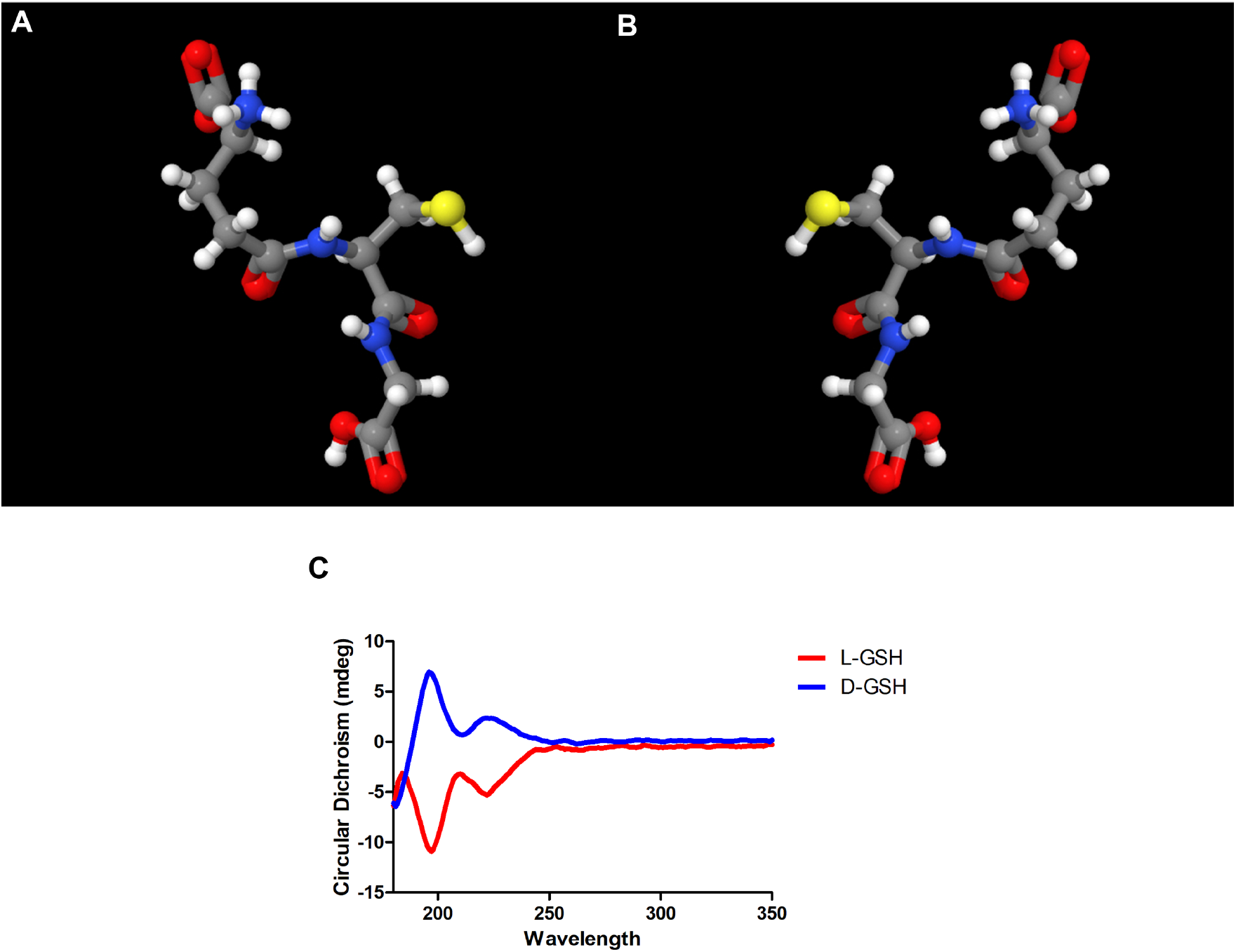
Characteristics of chiral GSHs. A, B Three-dimensional molecular structure of (A) L-GSH and (B) D-GSH. C CD spectra of L-GSH (red) and D-GSH (blue).

### D-GSH accelerated axon regrowth across the lesion center after SCI

To evaluate the neurological differences caused by the chirality of GSHs, we used a traumatic contusion SCI model for rats and a vehicle, L-GSH or D-GSH, as a treatment at the injured spinal cord. Afterwards, we examined axon regrowth using an anterograde tract-tracer (Biotinylated dextran amine; BDA). Axons in the vehicle group were mostly not seen past the lesion center (Figs 2a, d, and e). Axons in the L-GSH group also were mostly not seen past the lesion center (Figs 2b, f, and g). However, axons in the D-GSH group were observed past the lesion center (Figs 2c, h, and i). Specifically, total axon regrowth across the lesion center in the D-GSH group was over four-fold greater than that in the vehicle or L-GSH group (Fig EV1, Vehicle group: 1.95 ± 0.85, L-GSH group: 2.28 ± 1.69, D-GSH group: 8.62 ± 0.43, ^***^*p* < 0.001).

**Figure 2.**
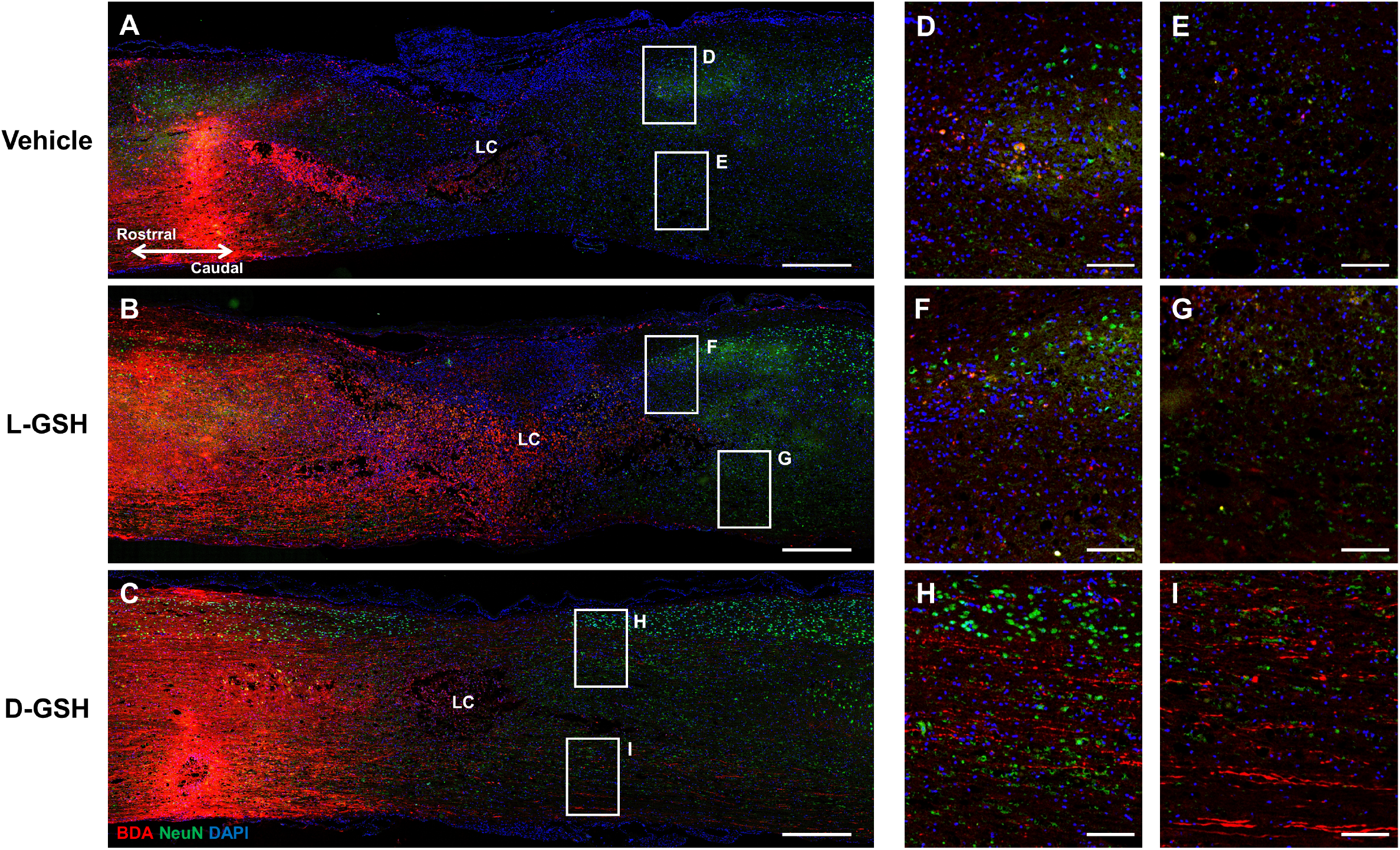
Axon regeneration across lesion center. BDA-labelled axons in composite tiled scans of sagittal sections stained for neuron (anti-NeuN, Green), axon (anti-streptavidin, Red), and cell nuclei (DAPI, Blue). LC, lesion center. A-C Representative images of the (A) vehicle group, (B) L-GSH group, and (C) D-GSH group after SCI in rats. Scale bar = 500 μm. D, E Higher magnification of boxed areas past the lesion center from (A). Scale bar = 100 μm. F, G Higher magnification of boxed areas past the lesion center from (B). Scale bar = 100 μm. H, I Higher magnification of boxed areas past the lesion center from (C). Scale bar = 100 μm.

Severed axons in the injured spinal cord do not transmit nerve impulses from the brain. Our results showed that axon regrowth was greater in the D-GSH group than in the L-GSH, indicating that the axon regeneration ability differed significantly according to the chirality of GSH.

### D-GSH reduced secondary damage after SCI and improved locomotor function

To investigate potential molecular mechanisms underlying this axon regrowth, we examined secondary damage after SCI in rats. As shown in Fig 3a, the lesion center in the vehicle group indicated the presence of expanded cavity spaces. The lesion center in the L-GSH group was also observed in the cavity spaces. However, cystic cavities in the D-GSH group were markedly decreased around the lesion center. The quantified tissue volume was increased in the D-GSH group compared to that in the vehicle group (Fig 3b, Vehicle group: 62.69 ± 6.58, D-GSH group: 82.25 ± 0.40, ^***^*p* < 0.001). In particular, the tissue volume in the D-GSH group was significantly increased compared to that in the L-GSH group (L-GSH group: 65.92 ± 2.74, ^**^*p* < 0.01).

**Figure 3.**
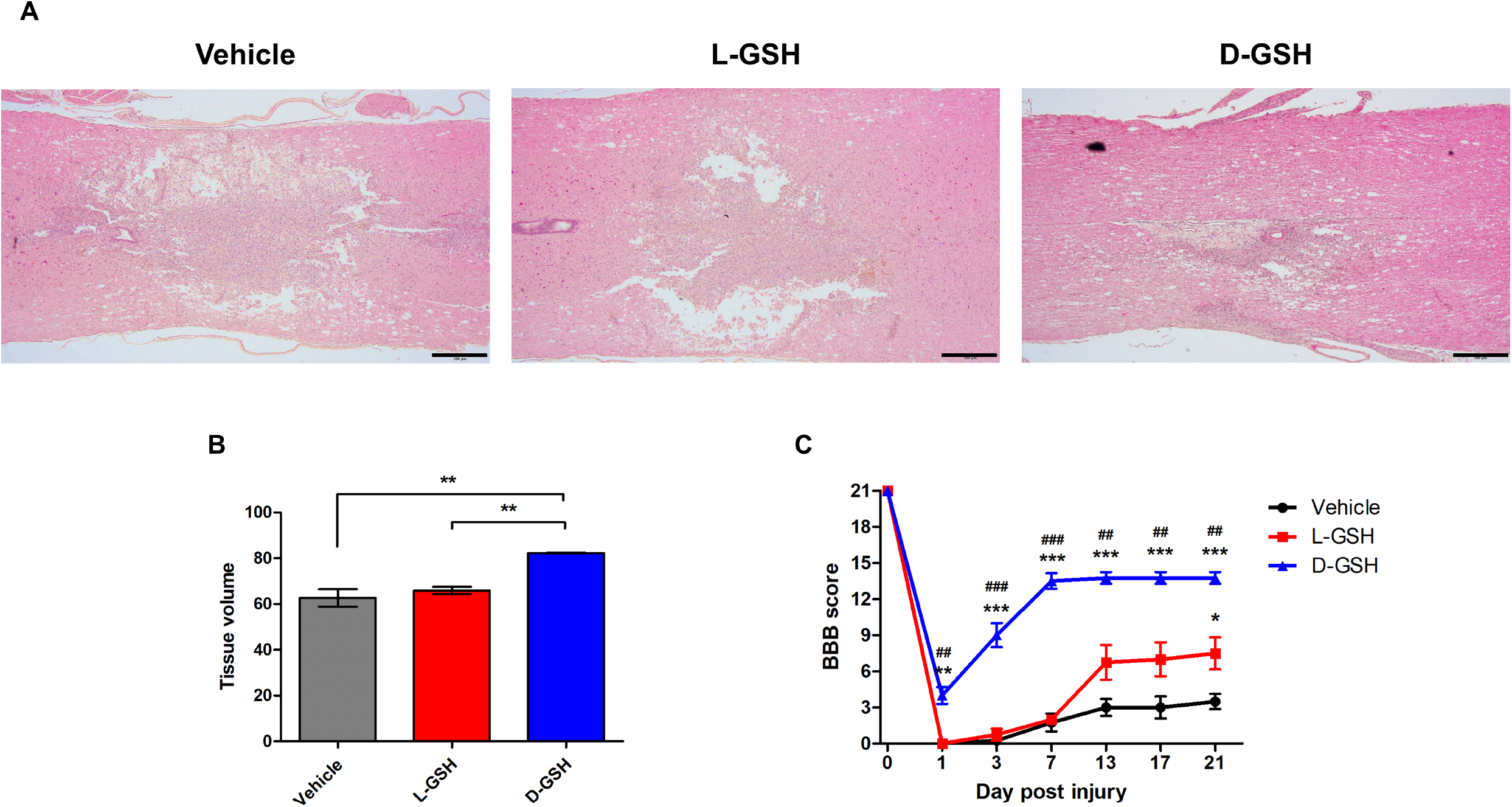
Histology and hindlimb locomotion. Histopathological changes and behavior functions were evaluated for the vehicle-, L-GSH-, and D-GSH-treated rats. A Representative images of the spinal cord stained with H&E in the vehicle, L-GSH, and D-GSH groups. B Quantitative analysis of the tissue volume from (A). C Comparison of BBB locomotor scores in the vehicle, L-GSH, and D-GSH groups. The BBB scores were evaluated for 21 days after SCI. ^*^ denotes a significant difference between the vehicle group and the D-GSH group. ^#^ denotes a significant difference between the L-GSH group and the D-GSH group. Results are the mean ± standard deviation (SD); ^*^p < 0.05, ^**^p and ^##^p < 0.01, and ^***^p and ^###^p < 0.001

Afterwards, we evaluated whether increasing axon regrowth could essentially improve functional recovery after SCI in rats (Fig 3c). The motor function was assessed on 1, 3, 7, 13, 17, and 21 days post-injury (DPI) using the Basso, Beattie, and Bresnahan (BBB) hindlimb locomotor rating scale (Basso *et al*, 1995). On 1 DPI, there were no differences in the extent of locomotor dysfunction in the vehicle or L-GSH group. However, the locomotor score of the D-GSH group was significantly increased compared to those of the vehicle and L-GSH group (D-GSH group: 4.00 ± 1.41, ^**^*p* < 0.01, ^##^*p* < 0.01). On 3 and 7 DPI, motor functions of D-GSH group were also significantly improved compared to those of the vehicle and L-GSH group (3 DPI: Vehicle group: 0.25 ± 0.50, L-GSH group: 0.75 ± 0.96, D-GSH group: 9.00 ± 2.00; 7 DPI: Vehicle group: 1.75 ± 1.50, L-GSH group: 2.00, D-GSH group: 13.50 ± 1.29, ^***^*p* < 0.001, ^###^*p* < 0.001). The locomotor score of the D-GSH group reached a plateau on 7 DPI at 13 points when the locomotor scores of the vehicle group and the L-GSH group reached a plateau on 13 DPI at 3 points and 7 points, respectively.

Macaya and Spector reported that the increased tissue repairing around cavity spaces induced axon regrowth (Macaya & Spector, 2012). Moreover, Hong et al. found that functional recovery was accompanied by the preservation of motor neurons and increased axon regeneration (Hong *et al*, 2017). Our findings showed that D-GSH limited the spread of cystic cavities and improved functional recovery more compared to L-GSH, demonstrating that D-GSH induces the regrowth of dissected axons by suppressing secondary damage.

### D-GSH inhibited glial scar formation after SCI

After SCI, the excessive inflammatory response induces glial scar formation, which can cause permanent paraplegia by disturbing axon contact (Ren *et al*, 2018). We investigated whether D-GSH provided stronger inhibition of the inflammatory response and glial scar formation than L-GSH (Fig 4). Tumor necrosis factor-α (TNF-α) is a polypeptide cytokine that plays a major role in the immune and inflammatory activities of the central nervous system (CNS). TNF-α fluorescence intensity levels in the vehicle group and L-GSH group were detected at 14.72 ± 1.47 and 5.04 ± 2.45, respectively. However, the TNF-α fluorescence intensity in the D-GSH group was significantly decreased compared to that in the vehicle group (D-GSH group: 0.32 ± 0.19, ^**^*p* < 0.01). In addition, the TNF-α fluorescence intensity in the D-GSH group was significantly decreased compared to that in the L-GSH group (^*^*p* < 0.05). Astrocytes, the major glial cell type in the CNS, are important contributors to inflammatory immune responses. They produce several neurotrophic substances that regulate the viability of neurons after injury, but they are also a source of pro-inflammatory cytokines such as TNF-α, interleukin (IL)-1β, and IL-6 (Stoll *et al*, 1998). Activated astrocytes produce a glial fibrillary acidic protein (GFAP) and form a glial scar, which blocks axon regrowth in the CNS. The GFAP fluorescence intensities in the vehicle group and L-GSH group were detected at 14.76 ± 3.47 and 11.44 ± 1.22, respectively. However, the GFAP fluorescence intensity in the D-GSH group showed a significant decrease compared to that in the vehicle group (D-GSH group: 4.16 ± 1.61, ^**^p < 0.01). Taken together, these results indicate that D-GSH provides stronger suppression of glial scar by decreasing the production of TNF-α and GFAP compared to L-GSH.

**Figure 4.**
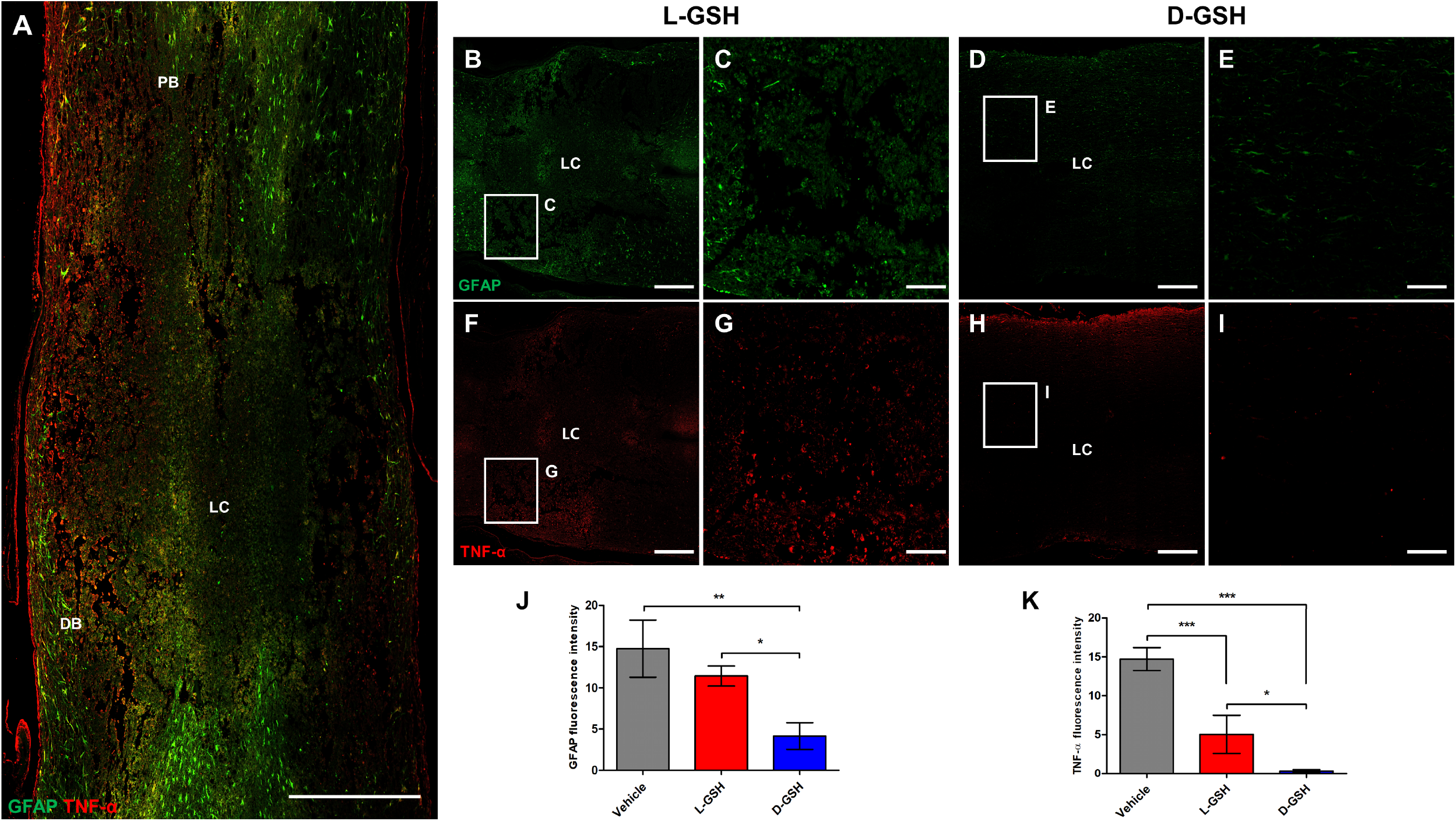
Glial scar formation. Immunofluorescence analysis in composite tiled scans of sagittal sections stained for glial scar (anti-GFAP, Green), inflammatory cytokine (anti-TNF-α, Red). PB, proximal border; DB, distal border. A Representative merged image for GFAP and TNF-α in the vehicle group. Scale bar = 500 μm. B Representative image for GFAP in the L-GSH group. Scale bar = 500 μm. C Higher magnification of the proximal border (boxed area) from (B). Scale bar = 100 μm. D Representative image for GFAP in the D-GSH group. Scale bar = 500 μm. E Higher magnification of the proximal border (boxed area) from (D). Scale bar = 100 μm. F Representative image for TNF-α in the L-GSH group. Scale bar = 500 μm. G Higher magnification of the proximal border (boxed area) from (F). Scale bar = 100 μm. H Representative image for TNF-α in the D-GSH group. Scale bar = 500 μm. I Higher magnification of the proximal border (boxed area) from (H). Scale bar = 100 μm. J, K Quantitative analyses of the fluorescence intensity for (J) GFAP and (J) TNF-α. Results are the mean ± standard error of the mean (SEM); ^*^*p* < 0.05, ^**^*p* < 0.01, and ^***^*p* < 0.001.

### Chiral GSH decreased the production of inflammatory cytokines after SCI

For a more in-depth comparison of the anti-inflammatory effect of the chiral GSHs, we measured inflammatory cytokines in injured spinal cords of vehicle-, L-GSH, and D-GSH-treated rats. The secretion levels of TNF-α and IL-1β in the L-GSH group were significantly lower than those in the vehicle group (Fig 5a, ^*^*p* < 0.05). However, the secretion levels of IL-6 and cyclooxygenase-2 (COX-2) in the L-GSH group were not significantly decreased compared to those in the vehicle group. On the other hand, the secretion level of TNF-α in the D-GSH group was significantly suppressed compared to in the L-GSH group (^*^*p* < 0.05). Moreover, this tendency was also found in other inflammatory cytokines, in this case IL-1β, IL-6 and COX-2 (Figs 5b-d, ^**^*p* < 0.01, ^***^*p* < 0.001).

**Figure 5.**
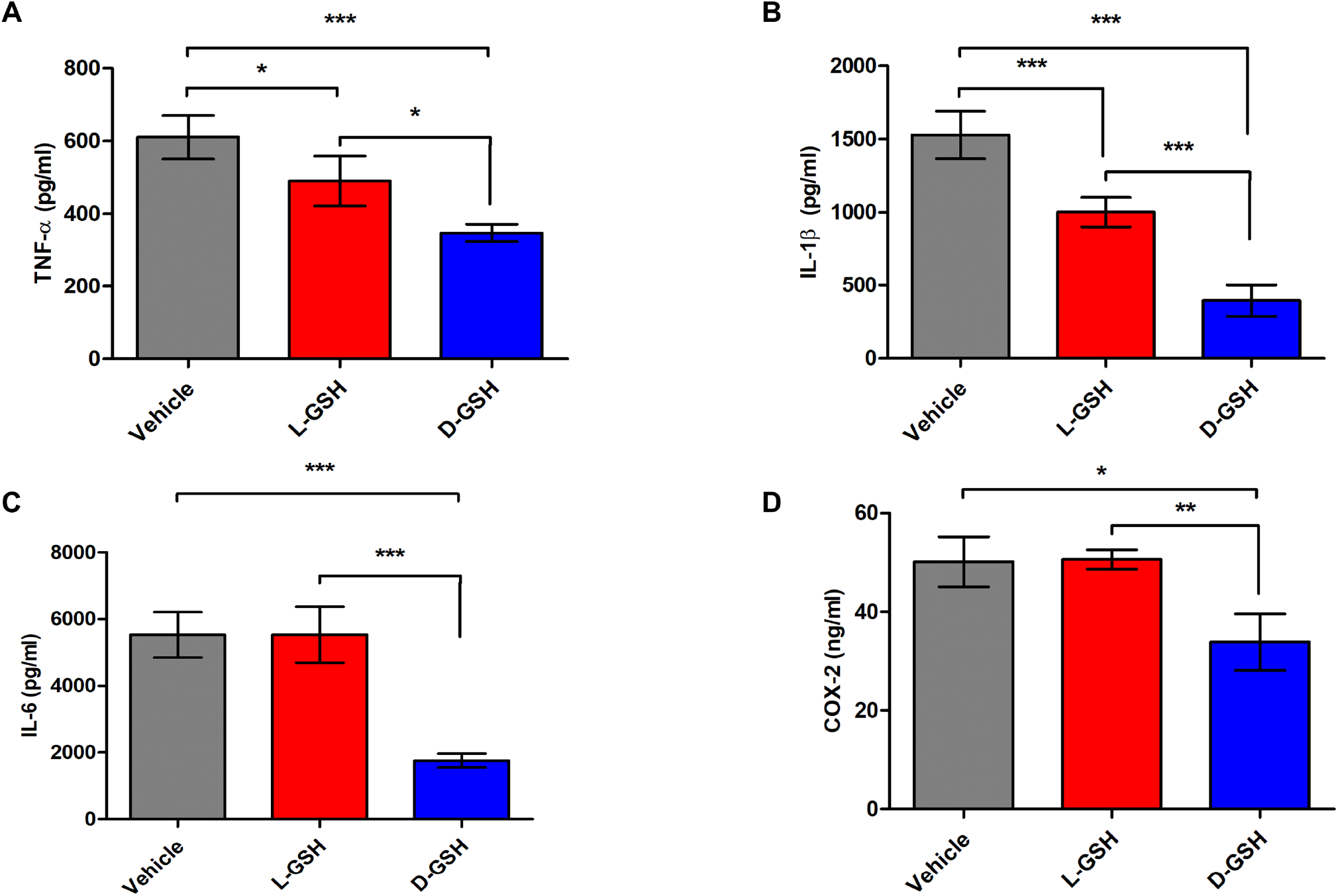
Production of inflammatory cytokines after SCI. A-D Secretion levels of inflammatory cytokines in the vehicle, L-GSH, and D-GSH groups after SCI. ELISA quantitation of (A) TNF-α, (B) IL-1β, (C) IL-6, and (D) COX-2. Results are the mean ± SEM; ^*^*p* < 0.05, ^**^*p* < 0.01, and ^***^*p* < 0.001.

In an injured spinal cord, macrophages are differentiated into M1 or M2 macrophages. The number of polarized M1 macrophages peaks at one week after injury and exacerbates neuroinflammation by secreting inflammatory cytokines such as TNF-α and IL-6 (Ko *et al*, 2017). On the other hand, polarized M2 macrophages induce anti-inflammatory cytokines and growth factors (Kim *et al*, 2018). As shown in Fig 5, D-GSH significantly decreased inflammatory cytokines. This result indicates that the differentiation into M1 macrophages was inhibited by D-GSH in SCI rats. Therefore, this study demonstrates that D-GSH decreases the inflammatory response by suppressing differentiation into M1 macrophages more compared to that by L-GSH.

### Chiral GSH suppressed inflammation through the mitogen-activated protein kinase (MAPK) signaling pathway after SCI

To elucidate the anti-inflammatory pathway of the chiral GSHs, we examined the activation of the MAPK pathway by western blotting (Figs 6a-d). The phosphorylated forms of the extracellular signal-regulated kinase (ERK), c-Jun N-terminal kinase (JNK), and p38 signals in the MAPK pathway are the principle processes that function during the inflammatory response. The fold ratio of the vehicle group was set to 1-fold and the relative fold change was calculated (Figs 6e-g). The p/t form volume of ERK in the L-GSH group was not lower compared to that in the vehicle group (L-GSH group: 0.87 ± 0.09). However, the p/t volume of ERK in the D-GSH group was decreased as compared to that in the vehicle group (D-GSH group: 0.67 ± 0.13, ^*^*p* < 0.05). The p/t volume of JNK in the L-GSH group was decreased as compared to that in the vehicle group (L-GSH group: 0.81 ± 0.03, ^**^*p* < 0.01). However, the p/t volume of JNK in the D-GSH group was decreased as compared to those in the vehicle and L-GSH groups (D-GSH group: 0.55 ± 0.01, ^**^*p* < 0.01, ^***^*p* < 0.001). Interestingly, the p/t volume of p38 in the L-GSH group was increased as compared to that in the vehicle group (L-GSH group: 1.18 ± 0.01, ^*^*p* < 0.05). On the other hand, the p/t volume of p38 in the D-GSH group was also decreased as compared to those in the vehicle and L-GSH groups (D-GSH group: 0.81 ± 0.10, ^*^*p* < 0.05, ^**^*p* < 0.01).

**Figure 6.**
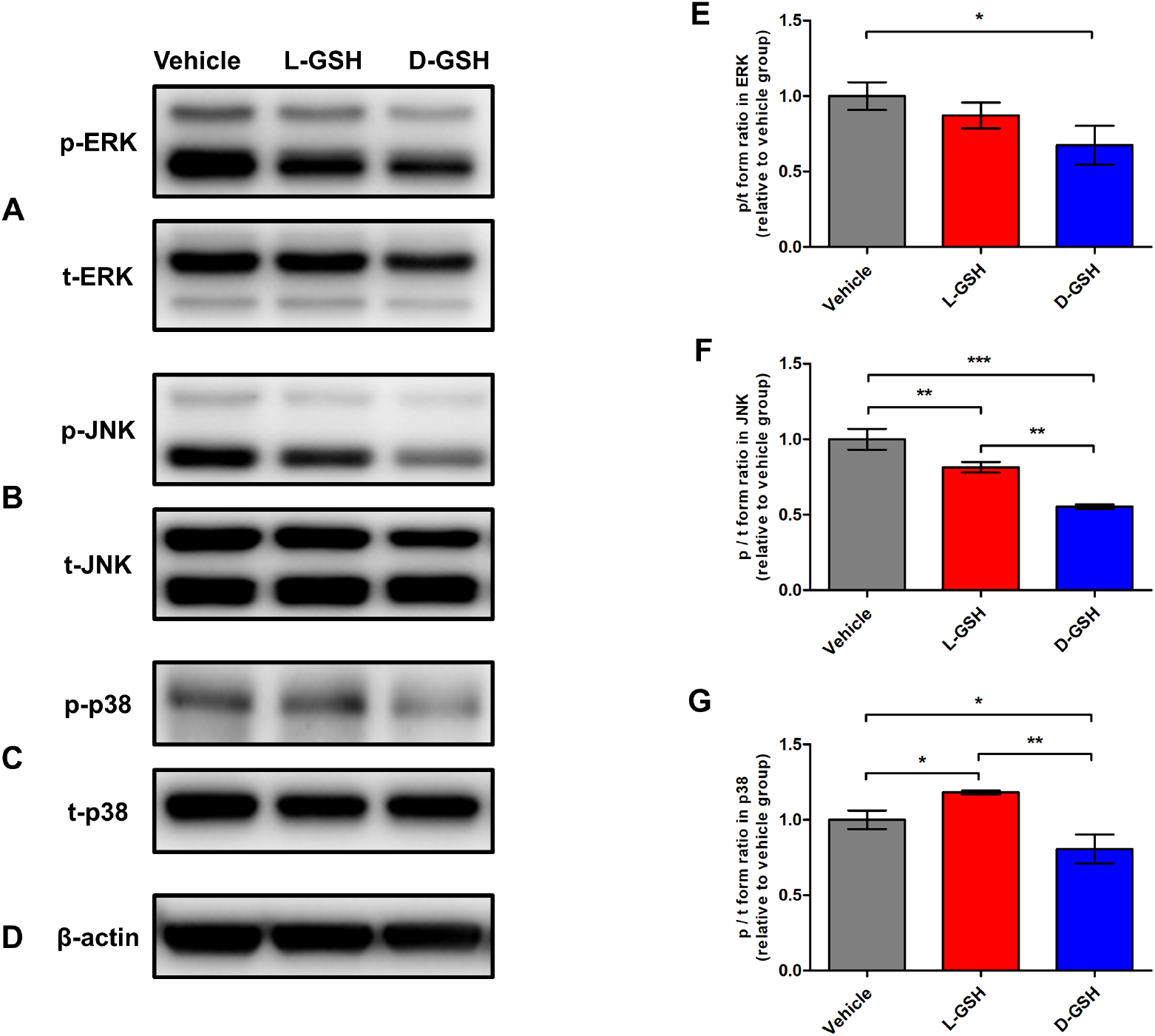
Western blots. The phosphorylation activities of the MAPK signaling pathway in the vehicle, L-GSH, and D-GSH groups. A-D Representative images of the p and t forms of (A) ERK, (B) JNK, (C) p38, and (D) β-actin. E-G Quantitative analyses of the p/t forms of (E) ERK, (F) JNK, and (G) p38. The p/t form volume in the vehicle group was set to 1-fold, and the ratio was relatively calculated and quantified. Results are the mean ± SEM; ^*^*p* < 0.05, ^**^*p* < 0.01, and ^***^*p* < 0.001.

Inflammatory responses are typically triggered by major intracellular signaling pathways of the MAPK pathway (Kyriakis & Avruch, 2001). Specifically, activation of the MAPK pathway increases the production of interferon and inflammatory cytokines (Cargnello & Roux, 2011), leading to the migration of macrophages (Sunderkotter *et al*, 1994). As shown in Fig 6, phosphorylation of ERK, JNK, and p38 in the D-GSH group decreased more than those in the L-GSH group. These findings emphasize the different anti-inflammatory effects of chiral GSHs.

### Different uptake levels of chiral GSHs in immune cells

To examine the anti-inflammatory mechanism of chiral GSH, we calculated the absorbance of various GSH concentrations using UV-visible spectroscopy. A declining tendency was observed according to an increase in the GSH concentration at 560 nm (Fig 7a). Afterwards, we treated lipopolysaccharide-stimulated bone marrow-derived macrophage (BMDM) cells with L- or D-GSH on 24 hours and analyzed the GSH uptake rates. The fold ratio of the L-GSH group was set to 1-fold and the relative fold change was calculated. Interestingly, the GSH uptake ratio in the D-GSH-treated cells was significantly higher than that in the L-GSH-treated cells (Fig 7b, D-GSH group: 1.51 ± 0.07, ^***^*p* < 0.001).

**Figure 7.**
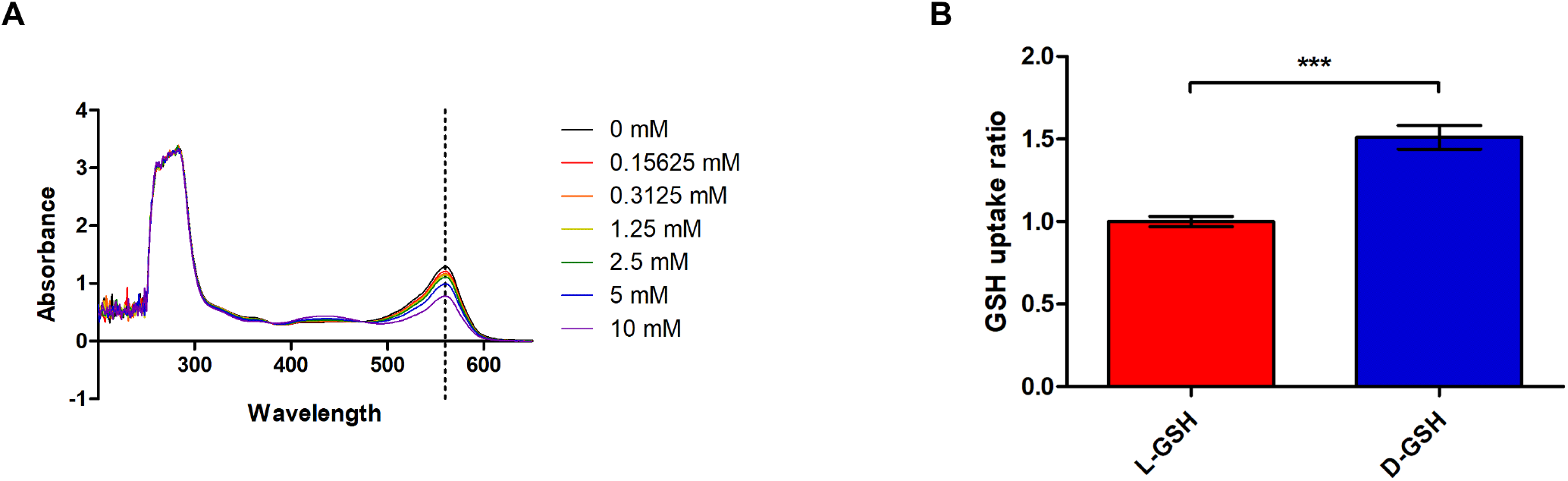
Intracellular uptake ratio. Comparative evaluation of the intracellular uptake ratios of the chiral GSHs A Absorbance of various GSH concentrations B Intracellular uptake ratios of macrophages treated with L- or D-GSH based on (A). The fold ratio of the L-GSH group was set to 1-fold and the relative fold change was calculated. Results are the mean ± SEM; ^*^p < 0.05, ^**^p < 0.01, and ^***^p < 0.001.

After SCI, nitric oxide (NO) radicals and superoxide radicals form peroxynitrite (Xiong *et al*, 2007), inducing vascular relaxation (Villa *et al*, 1994). This accelerates the pro-inflammatory response by increasing the infiltration of activated macrophages (Pacher *et al*, 2007). GSH plays a major role in maintaining redox homeostasis and the direct scavenging of NO radicals owing to its high concentration in cells (Jeong *et al*, 2018). Recently, Yeom et al. demonstrated that D-chirality had higher cellular uptake than L-chirality (Yeom *et al*., 2020). Here, we found that D-GSH showed a higher cellular uptake ratio than L-GSH. In light of those earlier results together, our study suggest that D-GSH can decrease the inflammatory response given its high uptake ability with activated macrophages more compared to L-GSH.

In this study, D-GSH increased axon regrowth past the lesion site and improved functional recovery by suppressing secondary damage. Furthermore, D-GSH inhibited inflammatory cytokines, suppressed the MAPK signaling pathway, and showed more interaction with activated macrophages than L-GSH. Although the anti-inflammatory potential of L-GSH has been reported for other anatomical sites (Ghezzi, 2011; Guizar-Sahagun *et al*., 2005; Villa *et al*, 2002), to the best of our knowledge, this is the first report to investigate chirality-dependent anti-inflammatory effects of chiral GSHs after SCI in an animal model. Our findings suggest that chiral GSH can serve as an alternative drug for SCI. It may also have therapeutic potential as a treatment for other diseases.

## Materials and methods

### Preparation and characterization of chiral GSHs

L-GSH was purchased from Sigma Aldrich (St. Louis, MO) and D-GSH was obtained from KOMA biotech (Seoul, South Korea). The chirality of the GSHs was measured by CD spectroscopy (Chirascan plus, Applied Photophysics, UK). For the in vivo experiment, L- or D-GSH was dissolved to a concentration of 10 mM using 0.9% sterile saline in each case.

### Surgical procedures

All rats were handled in accordance with the regulations of the Institutional Animal Care and Use Committee (IACUC) of CHA University (IACUC190162) and according to the Guide for the Care and Use of Laboratory Animals (NIH; Bethesda, MD, USA). Experiments were conducted on seven-month-old female Sprague Dawley rats (170 − 210 g body weight). Twenty-four rats were divided into a vehicle group (n = 8), an L-GSH group (n = 8) and a D-GSH group (n = 8).

For SCI, after the rats were anesthetized, a midline incision was made in the lower back. Tissues were dissected layer by layer to reveal the T8-T10 vertebra. A T9 total laminectomy was conducted to expose the dura. The spinous processes were fixed by clamps. A stainless-steel rod (2.0 mm diameter and 40 g weight) was used to apply a contusion injury. After the injury, the weight rod was quickly removed. Each treatment was applied stereotaxically onto the SCI lesions 1 mm below the surface at 1 uL per minute for five minutes. Each dose was repeated three times around the lesion center. After the injury, the surgical site was closed layer by layer. Tract-tracing of axons was performed via BDA injections (Sigma; #D1956). The BDA injections were performed one week after SCI to allow time for molecular expression. BDA was injected into two sites (one on each side of the cord, 0.5 μL (dissolved in sterile saline)) 1.5 mm below the surface at 0.1 μL per minute using a 33-gauge Hamilton syringe. After surgery, the rats were kept warm and housed separately, with free access to food. On each day at 8 am and 8 pm, a bladder massage was conducted to assist with urination. All surgeries were performed by the same spine neurosurgeon (S. Sohn).

### Hindlimb locomotor evaluation

To test the hindlimb function, open-field locomotion was evaluated using BBB scores (Basso *et al*., 1995). After SCI, the rats were evaluated on 1, 3, 7, 13, 17, and 21 DPI. Two trained investigators who were blind to the experimental conditions performed the behavioral analyses.

### Histology and immunohistochemistry

Seven days after SCI, three rats from each group were anesthetized and perfused transcardially with 4% paraformaldehyde. The spinal cords were processed for hematoxylin and eosin (H&E) staining and immunofluorescence, as described in the literature (Kim *et al*, 2019). Primary antibodies were as follows: rabbit anti-GFAP (1:1,000), mouse anti-TNF-α (1:1,000), and rabbit anti-NeuN (1:1,000) all from Abcam. Fluorescence secondary antibodies were conjugated to Alexa 488 or Alexa 568 (Molecular Probes). BDA tract-tracing was visualized with streptavidin-Alexa 594 (1:1,000, Thermofisher). Nuclei were stained with 4′,6′-diamidino-2-phenylindole dihydrochloride (DAPI; 2 ng/ml, Molecular Probes). The sections were mounted using a fluorescence mounting medium (DAKO, Glostrup, Denmark). Sections were examined and photographed using light microscopy (DP74, Olympus, Japan) or confocal laser scanning microscopy (LSM 880, Carl Zeiss, Germany).

### Quantification of inflammatory cytokines

Segments of the spinal cord (10 mm) including the lesion center were collected after SCI. The segments were collected and washed with phosphate-buffered saline and homogenized in lysis buffer. Afterwards, they were centrifuged at 13,000 rpm at 4 °C for 15 minutes. Protein concentrations in the tissue lysates were measured with the aid of a bicinchoninic acid protein assay kit (Thermo Scientific, Rockford, IL). TNF-α, IL-1β, IL-6 (Koma Biotech, Seoul, South Korea), and COX-2 (CUSA bio, Hubei, China) enzyme-linked immunosorbent assay (ELISA) kits were used according to each manufacturer’s directions.

### Western blotting

Equal amounts of protein (30 μg) were subjected to sodium dodecyl sulfate-polyacrylamide gel electrophoresis and transferred to nitrocellulose membranes. The membranes were incubated with 5% skim milk for one hour to block the nonspecific binding and probed with primary antibodies with the phosphorylated forms of the ERK (p-ERK; 1:1,000), JNK (p-JNK; 1:1,000), and p38 (p-p38; 1:1,000). Subsequently, equal membranes were stripped and reprobed with the total forms of ERK (t-ERK; 1:1,000), JNK (t-JNK; 1:1,000), and p38 (t-p38; 1:1,000). All primary antibodies were purchased from Cell Signaling Technology (Danvers, MA, USA), except for β-actin (1:5000, Abcam, Cambridge, UK). As an internal control, β-actin was probed into the membranes. The primary antibodies were followed by incubation with secondary antibodies (1:5,000; Santa Cruz Biotechnology, Dallas, TX). The visualized signal bands were detected using an ECL solution (Amersham, Buckinghamshire, UK) through a ChemiDoc XRS System (Bio-Rad, Hercules, USA). The p/t form volumes were calculated and quantified using ImageJ software. The p/t form volume of the vehicle group was set to 1-fold and the ratio change of the normalized fold was relatively calculated and quantified.

### Isolation of bone marrow-derived macrophages

The isolation protocol from rat bone marrow by Virginie et al. (Trouplin *et al*, 2013) was used. Briefly, rats were euthanized by CO_2_ asphyxiation and the femur and tibia were collected aseptically. Each bone was cut at the joint. Bone marrow was collected by rinsing the bone inner cavity with Dulbecco’s modified Eagle’s medium (DMEM) containing 10% fetal bovine serum (FBS), 2% glutamate, and 1% penicillin. The collected bone marrow was centrifugated (10 minutes, 450 × g). Erythrocytes were lysed via red blood lysis buffer (Sigma; #37757) for 30 seconds. Then, DMEM was quickly added to the cells, after which they were centrifuged. To eliminate resident macrophages, the cells were filtered with a 40 μm membrane and were incubated for four hours at 37 ℃ in tissue-culture-treated plates. Afterwards, supernatants were collected and centrifuged, the pellet was dissociated in a 150 ml of a medium consisting of complete DMEM (cDMEM) containing 10% FBS, 2% glutamate, 1% penicillin, and 20 ng/ml recombinant rat macrophage colony-stimulating factor (#315-02; Peprotech, NY, USA). Cells were distributed at 10 mL of suspension per petri dish (15 Petri dishes; Corning, #430591) and cultivated at 37 °C and 5% CO_2_. After three days, we added 10 mL of cDMEM to each petri dish. The dishes were incubated for four additional days. At seven days, the BMDM cells were harvested and seeded for the indicated experiments.

### Calculation of intracellular uptake rates

The absorbance levels of various GSH concentrations (0, 0.15625, 0.3125, 1.25, 2.5, 5, and 10 mM) were calculated by UV-visible spectroscopy. These values formulated the GSH standard curve. Then, BMDM cells were seeded on a petri dish at a density of 1 × 10^7^ cells/dish and treated with 10 mM L- or D-GSH for 24 hours. The absorbance of each supernatant was measured by a microplate reader (Bio-Rad, Hercules, USA) at a wavelength of 560 nm. Afterwards, we analyzed the uptake ratios based on the GSH standard curve. The fold ratio of the L-GSH group was set to 1-fold and the relative fold change was calculated.

### Statistical analyses

Analysis of variance (ANOVA) was used to compare differences in the BBB score at different time points. Two-group comparisons were conducted with Student’s t-tests. Differences in p-values for which ^*^*p* < 0.05, ^**^*p* < 0.01, and ^***^*p* < 0.001 were considered to be statistically significant.

## Acknowledgements

The authors are indebted to Robert Langer, Institute Professor, Massachusetts Institute of Technology for his pro bono editing of this manuscript. This work was supported by the Basic Science Research Program through the National Research Foundation of Korea (NRF) funded by the Ministry of Science and ICT (NRF-2020R1F1A1069875) and by a grant from the Korea Health Technology R&D Project through the Korea Health Industry Development Institute (KHIDI), funded by the Ministry of Health & Welfare, Republic of Korea (HI20C0173)

## Author Contributions

Seong Jun Kim and Seil Sohn conceived and directed the project. Seong Jun Kim designed the experiments. Seong Jun Kim, Wan-Kyu Ko, Gong Ho Han, and Daye Lee carried out the experiments. Yuhan Lee, Seung Hun Sheen, and Je Beom Hong conducted the data analysis and interpreted the results. Seong Jun Kim wrote edited the paper. All authors reviewed the manuscript.

---

**Figure EV1.**
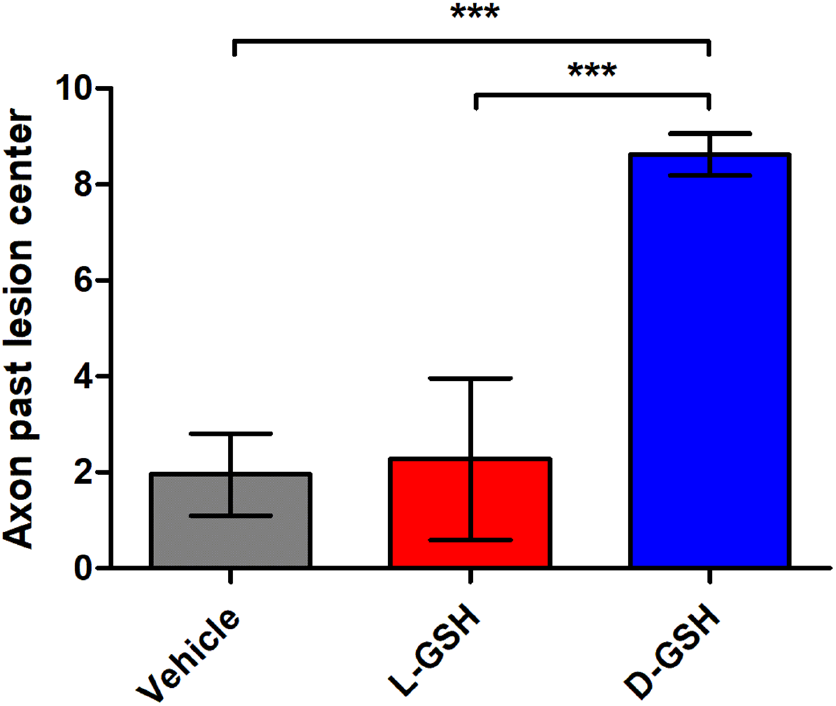
Quantitation of BDA-labelled axon regrowth past the lesion center. Quantitative analyses of BDA-labelled axons past the lesion center in the vehicle, L-GSH, and D-GSH groups are shown. Results are the mean ± SEM; ^*^*p* < 0.05, ^**^*p* < 0.01, and ^***^*p* < 0.001.

